# Maturation-dependent splicing alterations constrain *SYNGAP1* splice-switching therapy

**DOI:** 10.64898/2026.08.26.745682

**Authors:** J.A. Kamp, K.N. Wijnant, N. Maas, D. Gülyurt, M.J.M. Rieder, M.A. Jolfaei, C. Gontan, S.A. Kushner, Y. Elgersma, L.E.L.M. Vissers, N. Nadif Kasri, F.M.S. de Vrij

## Abstract

Haploinsufficiency in *SYNGAP1* causes a severe neurodevelopmental syndrome. SYNGAP1 protein is mainly detected in neuronal synapses. However, *SYNGAP1* RNA is more widely expressed and strongly regulated via alternative splicing: alternative 3’ splice site (A3SS) inclusion leads to non-productive transcripts that are degraded through nonsense-mediated decay.

Recently, splice-switching oligonucleotides (SSOs) that redirect *SYNGAP1* splicing to increase SYNGAP1 protein levels were developed. However, we hypothesized that during neuronal maturation, non-productive splicing may decrease to enhance functional transcripts in mature neurons. This would reduce the abundance of the SSO target transcript, limiting the potential for SSO treatment to increase neuronal SYNGAP1 expression.

Using neural differentiation of human induced pluripotent stem cells, we show that the A3SS transcript is abundant in neural progenitors, astrocytes, microglia and immature neurons, with minimal presence in mature neurons.

These data imply that SSOs targeting A3SS might lack therapeutic efficacy to rescue the neuronal phenotypes associated with *SYNGAP1* haploinsufficiency.

**Graphical abstract:** 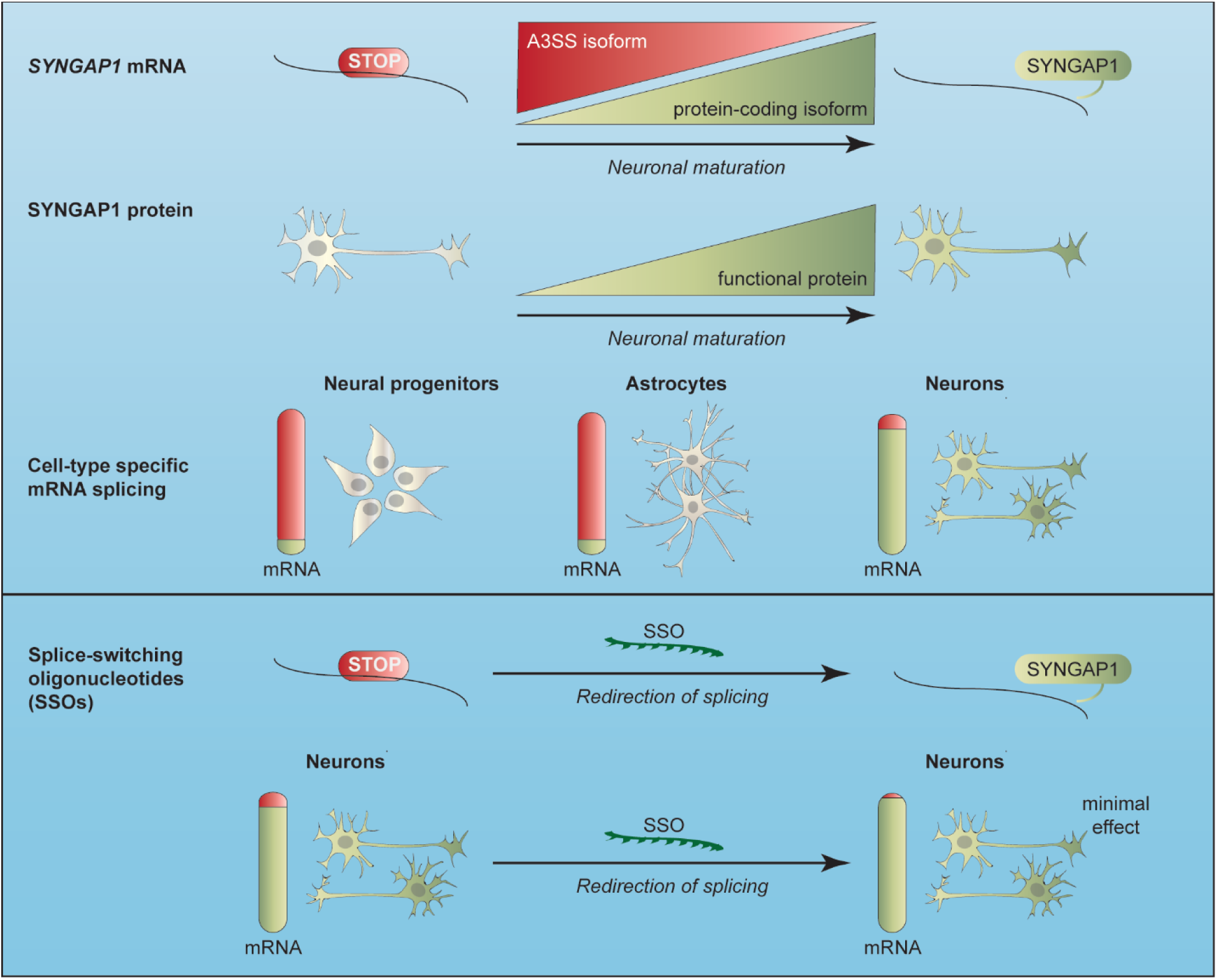

## Introduction

Synaptic plasticity, the adaptive modification of synapses through insertion or removal of postsynaptic receptors, underlies learning and memory. Mutations affecting the proteins involved in synaptic plasticity are key causes of learning deficits, such as intellectual disability. Mutations in the *SYNGAP1* gene, located on chromosome 6p21.32 and encoding Synaptic GTPase Activating Protein 1 (SYNGAP1), account for 0.5–1.0% of all intellectual disability cases, and lead to epilepsy and sensory processing impairments^1,2^. SYNGAP1 is an important regulator of long-term potentiation (LTP), the activity-dependent strengthening of synapses regarded as the neural basis of learning and memory^3,4^. In inactive synapses, SYNGAP1 prevents incorporation of α-amino-3-hydroxy-5-methyl 4-isoxazolepropionic acid receptors (AMPARs) into the postsynaptic membrane by competing with anchoring proteins^5^ and inhibiting small GTPases Ras and Rap^6^. This suppresses downstream signaling pathways, including the Ras-Raf-ERK-MEK pathway, that mediate production and incorporation of AMPARs^7–9^. During synaptic activity, phosphorylation of SYNGAP1 modulates its Ras- and Rap-inhibiting ability and causes its dispersion from the postsynaptic density (PSD) by reducing the affinity for PSD95, leading to increased AMPAR insertion and strengthening of the synapse^5,8,10–14^.

SYNGAP1 is a synaptic protein that is mainly detected in neurons. However, *SYNGAP1* RNA is present in non-neuronal brain cells and other tissues, but mainly in the form of an alternatively spliced isoform containing a premature stop codon. This isoform exists due to usage of an alternative 3’ splice site (A3SS) of exon 11. The A3SS alternatively spliced isoform is a substrate for nonsense-mediated decay. *SYNGAP1* alternative splicing works as a mechanism in non-neuronal cells to suppress excess SYNGAP1 expression^14^.

Recent research has focused on strategies to upregulate SYNGAP1 protein expression from the remaining functional allele to rescue phenotypes caused by *SYNGAP1* haploinsufficiency, including approaches based on TANGO (Targeted Augmentation of Nuclear Gene Output)^15^. The TANGO approach targets haploinsufficiencies using Splice-Switching Oligonucleotides (SSOs) to block non-productive pre-mRNA splicing, thereby forcing healthy alleles to increase the production of functional proteins in a mutation-independent manner. By redirecting the splicing of *SYNGAP1* pre-mRNA, researchers hypothesized that SYNGAP1 protein levels could be increased and compensate for the mutated allele that caused *SYNGAP1* haploinsufficiency^14,16^. SSOs that shift splicing of the alternative isoform to the functional isoform have been proposed as a therapeutic strategy, because they increase SYNGAP1 protein levels in neural cultures and cerebral organoids^14,16^. However, it was reported that A3SS inclusion rate in *SYNGAP1* mRNA is higher outside the brain and during early brain development compared to the adult brain in mice^14^. This raises the question whether A3SS-containing transcripts persist in mature neurons, where SYNGAP1 physiologically functions at excitatory synapses. The presence of A3SS inclusion in these neurons is a prerequisite for the therapeutic potential of SSOs, as they can only redirect splicing and functionally rescue neuronal phenotypes associated with *SYNGAP1* haploinsufficiency if the A3SS target transcript is available.

Because previous data suggest that the alternative isoform is mainly expressed in non-neuronal cells or during early neuronal development, we hypothesize that the observed increase in SYNGAP1 protein levels in organoids and neural cultures after splice switching primarily originates from astrocytes and progenitor cells in the cultures. To investigate this, we generated monocultures of human induced pluripotent stem cell (hiPSC)-derived neurons, astrocytes, and neural progenitors and additionally interrogated an independent RNA-sequencing resource that also includes hiPSC-derived microglia, and monitored alternative splicing of *SYNGAP1* exon 11. These data show that A3SS inclusion decreases during neuronal maturation, thereby diminishing the therapeutic potential of exon 11 SSOs for the treatment of *SYNGAP1* haploinsufficiency.

## Results

### SYNGAP1 protein expression is successfully restored after CRISPR/Cas9 mediated repair of a *SYNGAP1* mutant human induced Pluripotent Stem Cell (hiPSC) line and increases with neuronal maturation

To study *SYNGAP1*, we obtained a patient-derived hiPSC line harboring a heterozygous *de novo* loss-of-function mutation in *SYNGAP1* exon 17 (c.3718 C>T) and corrected the mutation using CRISPR/Cas9 editing to generate an isogenic control line (Figure ***1***A). To verify that SYNGAP1 protein levels are restored, we quantified protein levels in three- and five-week-old hiPSC-derived neuronal monocultures (Figure ***1***B). We found that SYNGAP1/GAPDH ratios significantly increased between weeks 3 and 5 in isogenic control neurons (mean±SD, week 3 = 0.543±0.049; mean±SD, week 5 = 1.027±0.174; p = 0.001; Figure ***1***C) and shows a trend towards increased SYNGAP1 levels in more mature mutant neurons (mean±SD, week 3 = 0.181±0.040; mean±SD, week 5 = 0.412±0.060; 95% CI [-0.024, 0.485], p = 0.076). Patient-derived neurons displayed significantly lower SYNGAP1/GAPDH ratios at both timepoints (week 3: p = 0.008; week 5: p < 0.001), indicating successful restoration of SYNGAP1 protein levels in isogenic control neurons.

**Figure 1.**
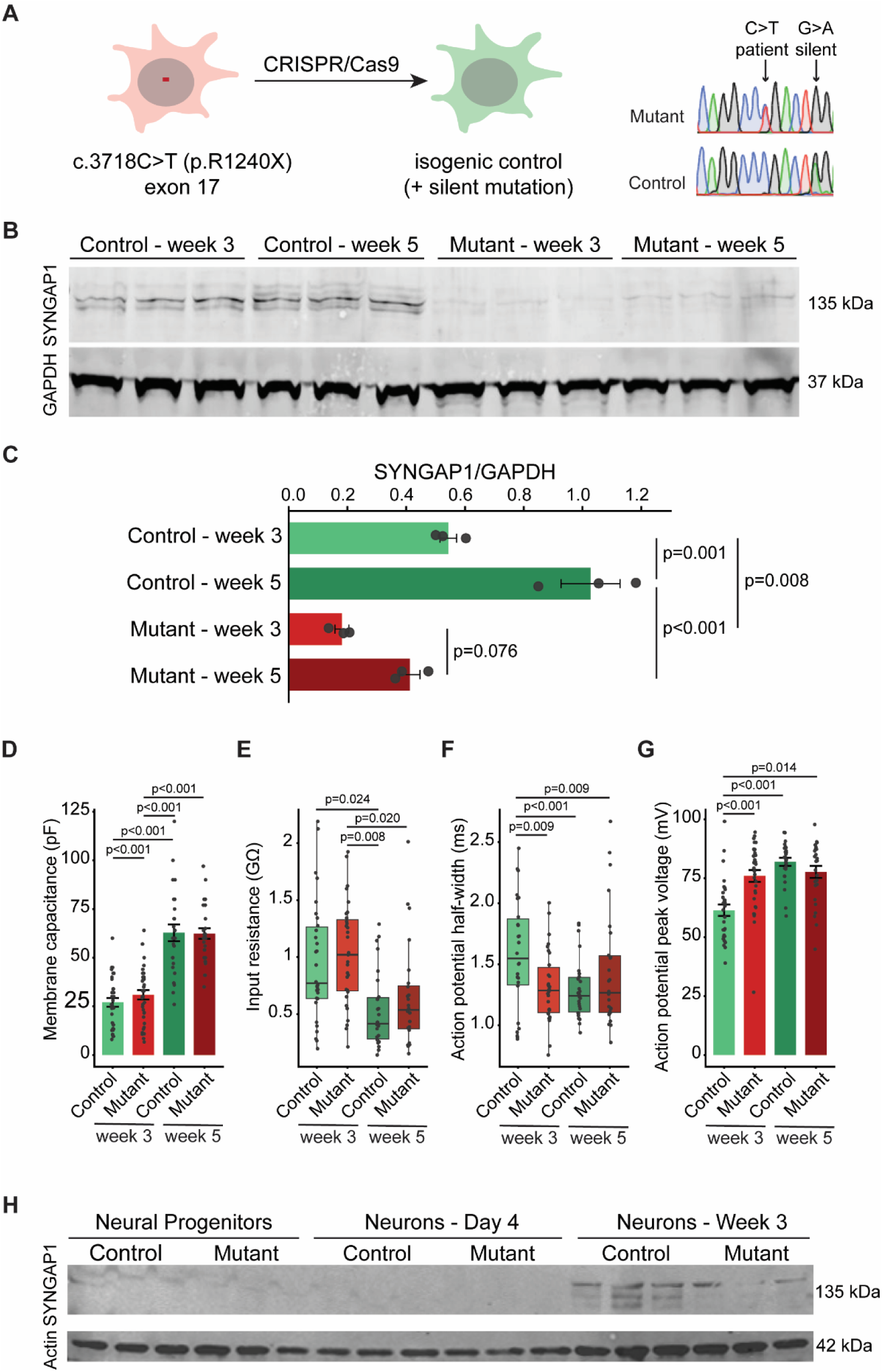
*SYNGAP1* haploinsufficient mature neurons show decreased protein levels and accelerated electrophysiological maturation compared to isogenic control. (A) Schematic representation of CRISPR/Cas9 editing and representative Sanger sequencing results of haploinsufficient hiPSCs and the generated isogenic control. (B) Western blot showing SYNGAP1 and GAPDH protein levels in control and mutant neurons in weeks 3 and 5. (C) Quantification of SYNGAP1 and GAPDH protein levels in control and mutant neurons in weeks 3 and 5. Dots represent measurements per replicate. Bars and error bars represent the mean and standard error, respectively. Ratios were compared using two-way ANOVA tests. (D-G) Electrophysiological properties of *SYNGAP1* haploinsufficient and isogenic control neurons at weeks 3 and 5. Dots represent individual neuronal measurements. Bar plots and error bars represent the mean and standard error. In boxplots, the center line, boxes, and whiskers represent the median, interquartile range (IQR), and minimum and maximum values within 1.5*IQR of the first and third quartiles, respectively. False Discovery Rate-corrected p-values were determined using linear mixed models controlling for batch differences. (D) Quantification of membrane capacitance. Control week 3: n=34; mutant week 3: n=36; control week 5: n=26; mutant week 5: n=31. (E) Quantification of input resistance. Control week 3: n=34; mutant week 3: n=36; control week 5: n=26; mutant week 5: n=31.(F) Quantification of action potential half-width. Control week 3: n=26; mutant week 3: n=30. Control week 5: n=26; mutant week 5: n=26. (G) Quantification of action potential peak voltage. Control week 3: n=30; mutant week 3: n=32 Control week 5: n=26; mutant week 5: n=26. (H) Western blot showing SYNGAP1 and Actin protein levels in control and mutant, neural progenitors and neurons at day 4 and week 3.

### *SYNGAP1* haploinsufficient Neurogenin (NGN2) neurons show earlier maturation of action potential properties

In *SYNGAP1* haploinsufficiency, higher basal incorporation of AMPARs in the postsynaptic membrane and limited LTP-dependent increases correspond to LTP defects and earlier maturation of excitatory synapses^11^. These alterations are consistently observed in both mouse ^17,18^ and human models of *SYNGAP1* haploinsufficiency, including patient-derived cortical organoids^19^, human embryonic stem cell (hESC)-derived cortical neurons transplanted into mice ^20^, and Neurogenin-2 (NGN2) neurons^4^. To verify that our NGN2 models are representative, we conducted patch clamping experiments to determine the electrophysiological maturation of our cultures.

Electrophysiological recordings were performed in 34 control and 36 patient-derived neurons in week (W) 3 and 26 control and 31 mutant neurons in week 5. There were no differences in passive properties (membrane capacitance, membrane resistance, input resistance or resting membrane potential) between genotypes at the same developmental timepoint, but maturation over time was observed as expected. For example, membrane capacitance significantly increased over time in both control neurons (mean±SD, W3 = 27.06±13.00 pF; mean±SD, W5 = 62.81±22.05 pF; β±SE = -25.81±4.02, p<0.001) and mutant neurons (mean±SD, W3 = 30.93±14.53 pF; mean±SD, W5 = 62.39±15.17 pF; β±SE = -23.25±3.70, p<0.001; Figure ***1***D). Input resistance significantly decreased between week 3 and week 5 in both control and mutant neurons (median (IQR) control W3 = 769.64 (629.08) MΩ; control W5 = 415.48 (364.10) MΩ; β±SE = 0.60±0.22, p=0.024; median (IQR) mutant W3 = 1020.16 (625.53) MΩ; mutant W5= 536.54 (376.85) MΩ; β±SE = 0.56±0.20, p=0.020, Figure ***1***E).

We found that action potential half-width was significantly shorter in mutant neurons in week 3 compared to control neurons of the same age (median (IQR) control = 1.54 (0.54) ms; mutant = 1.28 (0.37) ms; β±SE = 0.185±0.063, p=0.009, Figure ***1***F). This difference disappeared in week 5 (median (IQR) control = 1.24 (0.28) ms; mutant = 1.27 (0.46) ms; β±SE = -0.077±0.065, p=0.290; Figure ***1***F). Action potential half-width decreased over time in control (β±SE = 0.283±0.071, p<0.001), while no significant differences were observed in haploinsufficient neurons (β±SE = 0.021±0.067, p=0.754, Figure ***1***F). Together, these findings suggest *SYNGAP1* haploinsufficient neurons exhibit earlier maturation of action potential halfwidth compared to control neurons. Shorter action potential durations were due to a shorter rise time of week 3 mutant neurons compared to week 3 control neurons (median (IQR) control = 0.56 (0.22) ms; median (IQR) mutant = 0.38 (0.11) ms; p=0.023; Supplemental table S1). Similar to half-width, rise time was no longer significantly altered in week 5 (median (IQR) control= 0.36 (0.08) ms; median (IQR) mutant = 0.39 (0.14) ms; p=0.508). Rise time decreased over time in control neurons (p<0.001), while no significant differences were observed in haploinsufficient neurons (p=1). In addition, week 3 haploinsufficient neurons displayed a significantly higher action potential peak voltage than week 3 control neurons (mean±SD control = 62.39±17.90 mV; mutant = 80.14±17.81 mV; β±SE = -12.72±2.90, p<0.001, Figure ***1***G). No significant differences were found in week 5 (mean±SD control = 81.98±8.65 mV; mean±SD mutant = 77.70±12.99 mV; β±SE = 4.280±3.13, p=0.263; Figure ***1***G). A full overview of the electrophysiological properties of control and *SYNGAP1* mutant NGN2 neurons at weeks 3 and 5 can be found in supplemental table S1. These findings suggest that *SYNGAP1* haploinsufficiency accelerates maturation of action potentials, consistent with findings of accelerated maturation in other human and mouse models^17–21^.

### SYNGAP1 protein is not expressed during early stages of differentiation

SSO activity was previously investigated in young NGN2 neurons, at 4 days *in vitro* (DIV4)^14^. We wondered if SYNGAP1 is already expressed in such an early stage of differentiation. Therefore, we differentiated neurons and isolated protein after 4 days and 3 weeks of differentiation. We found that, similar to neural progenitor cells, SYNGAP1 protein is not present in DIV4 NGN2 neurons (Figure ***1***H), neither in control or mutant neurons. We verified that neuronal differentiation did occur successfully by staining for MAP2, NeuN and doublecortin (Supplemental figure 1). Also, astrocytes expressed GFAP and s100beta, but were negative for the neuronal marker MAP2 (Supplemental figure 1). Similarly, NPCs were negative for MAP2 and expressed Nestin and SOX2 (Supplemental figure 1). These data suggest that DIV4 neurons, even though they express neuronal markers as MAP2 and NeuN, are not expressing SYNGAP1 protein yet, and probably reflect a transition state between stem cells and neurons.

### *SYNGAP1* splicing is shifted towards the protein-coding isoform during maturation

Because we did not observe any SYNGAP1 protein produced by neurons after four days of differentiation, whereas it was expressed after three weeks of differentiation, we wondered whether we would see a shift in *SYNGAP1* RNA splicing from the A3SS isoform to the protein-coding isoform. Therefore, we differentiated neurons and isolated RNA after four days or three weeks after differentiation. To visualize the RNA that is usually degraded by nonsense-mediated decay (NMD), we treated the cultures with cycloheximide (CHX) for five hours before harvesting. We observed that the A3SS isoform was highly present, slightly more than the protein-coding isoform in four-day-old neurons (Figure 2BC): the percentage spliced in (PSI) of A3SS was 61% for CHX-treated controls and 66% for CHX-treated mutants. The splicing shifted dramatically towards the protein-coding isoform in three-week-old neurons (Figure 2BC), with A3SS PSI of 22% for CHX-treated controls and 29% for CHX-treated mutants, showing that *SYNGAP1* splicing alters during neuronal maturation. In concordance with previous reports^14,16^, we observed a significant effect of NMD inhibition in control day 4 neurons (mean DMSO = 52.5; mean CHX = 61.4, p<0.001; Figure 2C) and mutant day 4 neurons (mean DMSO = 60.6; mean CHX = 66.0, p=0.005; Figure 2C), while we did not observe a significant difference in week 3 neurons (mean control DMSO = 14.2; mean control CHX = 21.8, p=0.358; mean mutant DMSO = 22.7; mean mutant CHX = 28.6, p=0.473; Figure 2C). To verify our results, we analyzed data from the NMD alternative splicing (NMD AS) database ^22^. This database contains RNA sequencing data of hiPSC-derived wildtype excitatory neurons, astrocytes, and microglia treated with or without CHX, allowing mapping of NMD AS events (Figure 2DE). Linear modeling showed that cell type (p<0.001) and treatment (p=0.031) affect PSI scores, but there is no interaction between treatment and cell type (p=0.707), showing the effect of CHX treatment is similar in different cell types. In line with our findings in mutant and isogenic control neurons, wildtype DIV49 neurons exhibit a small increase in *SYNGAP1* alternative splicing after CHX treatment (Figure 2D), while it has been previously shown that CHX treatment had a stronger effect on NMD of different transcripts, like *NR4A2* and *CHD2*^22^. These data suggest that in relatively mature neurons, the rate of A3SS inclusion is low.

**Figure 2.**
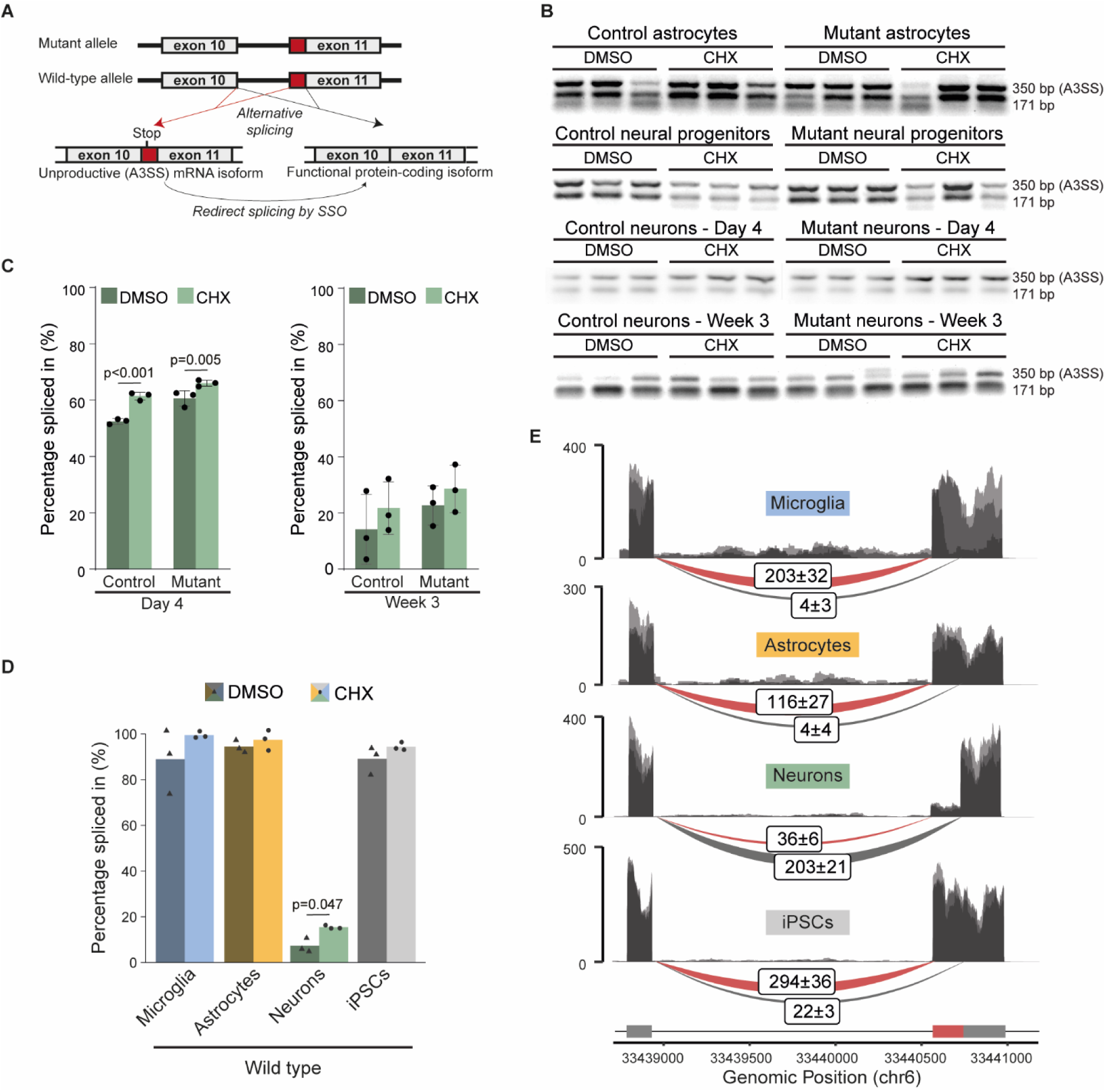
Differences in splicing of *SYNGAP1* mRNA in different iPSC-derived neural cell types and during neuronal maturation. (A) Alternative splicing of *SYNGAP1* leads to inclusion of a premature stop codon. Splice switching oligos can redirect splicing from the unproductive isoform to a functional protein-coding isoform. (B) RT-PCR results from neural progenitors, astrocytes, immature and mature excitatory neurons derived from patient and isogenic control iPSCs. Cells were treated with CHX or DMSO (n=3). (C) Percentage of A3SS inclusion in control and mutant iPSC-derived immature (left) and mature (right) excitatory neurons as determined by RT-PCR. Cells were treated with CHX or DMSO (n=3). Dots represent measurements per replicate. Means were compared using two-way ANOVA, p-values are False Discovery Rate-corrected. (D) Percentage of A3SS inclusion in wild-type iPSCs and iPSC-derived microglia, astrocytes, DIV 49 excitatory neurons treated with cycloheximide (CHX) or vehicle (DMSO) as determined by RNA sequencing (n=3), adjusted p-value determined by t-test. (E) Sashimi plots of control iPSCs and iPSC-derived microglia, astrocytes, mature excitatory neurons. The A3SS exclusion (grey) and inclusion (red) read numbers are indicated.

### The A3SS isoform is abundant in non-neuronal cells

We wondered whether other brain cell types also express the A3SS isoform, which would indicate that they could also be affected by SSO treatment. We generated astrocytes and neural progenitors, which do not produce SYNGAP1 protein (Figure 1H), and we observed clear presence of the A3SS isoform in these cell types, with and without NMD inhibition using CHX (Figure 2B), suggesting that SSO treatment may also alter splicing in neural progenitors and astrocytes. The finding that A3SS is clearly present in astrocytes is supported by data from the NMD alternative splicing (NMD AS) database^22^. These data show that in astrocytes, as well as in microglia and hiPSCs, the A3SS isoform is the predominant *SYNGAP1* isoform. This could explain the previously described effect of SSO treatment on SYNGAP1 protein levels in mature neural cultures and organoids^14,16^, which contain non-neuronal cell types like astrocytes and neural progenitor cells. The increased SYNGAP1 protein production might originate from astrocytes and neural progenitors, instead of the neurons, the cell-type of interest for treatment of *SYNGAP1* haploinsufficiency.

## Discussion

We showed that the primary target for SSO therapy of *SYNGAP1* haploinsufficiency, the A3SS isoform of *SYNGAP1* mRNA, decreases during neuronal maturation. Thus, in cells where SYNGAP1 protein is expressed, levels of the A3SS isoform are low. In contrast, other brain cells like astrocytes and microglia do show the A3SS isoform as the predominant *SYNGAP1* splice isoform. This has important implications for the treatment of patients with *SYNGAP1* haploinsufficiency, since the SSO’s working mechanism is to switch splicing from the A3SS isoform to the functional protein-coding isoform. Because cognitive deficits associated with *SYNGAP1* haploinsufficiency arise from dysfunction of forebrain excitatory neurons^23^ and the average age at diagnosis is approximately 5 years of age^24^, therapies designed for the treatment of *SYNGAP1* haploinsufficiency should be specific to mature neurons.

Genetically deleting the possibility of A3SS formation did rescue *Syngap1* haploinsufficiency-related phenotypes in mice^14^. However, in contrast to SSO treatment, the splice switching is already present at the start of brain development in this model. *SYNGAP1* is already expressed in radial glia during early brain development, and therefore *SYNGAP1* haploinsufficiency leads to disturbed cortical neurogenesis^25^. The early role of SYNGAP1 explains why not all behavioral phenotypes in *Syngap1* haploinsufficient mice can be improved by *Syngap1* gene restoration in adult mice^26^. However, because *SYNGAP1* haploinsufficiency diagnosis usually occurs postnatally, therapies should be tested on relatively mature brain models.

Testing therapies on mature human neurons poses a challenge. Neurons derived from hiPSCs can be used to overcome this challenge. However, at this moment, a consensus on maturation assessment is lacking. For example, maturation of the hiPSC-derived cells can be estimated by assessing functional maturity or by comparing the transcriptome with reference data of brain tissue at different developmental stages. Moreover, maturation of neurons differs greatly between differentiation protocols. Also between NGN2 overexpression protocols, maturity estimations of the resulting neurons vary remarkably. For example, it was proposed that week 6 NGN2 neurons resemble postnatal day-86 neurons *in vivo*^27^, while others find that week 7 NGN2 neurons largely reflect prenatal stages, with some molecular features of the postnatal brain^28^. Moreover, whether astrocytes are added to the neuronal culture greatly affects the functional maturation of the neurons, with increased maturation after addition of hiPSC-derived astrocytes versus rat astrocytes^29^. While giving precise comparisons of maturation of hiPSC-derived neurons compared to *in vivo* human neurons remains challenging, it is clear that NGN2 neurons do not reach the maturity of human neurons in a five year old brain, when patients suffering from *SYNGAP1* haploinsufficiency get diagnosed on average.

Our work emphasizes the importance of taking the maturation and cell-type diversity of hiPSC-derived neural models into account when developing therapies for neurodevelopmental disorders, especially when opting for the TANGO approach, since maturation and cell identity affect the availability of TANGO target transcripts. More knowledge on splicing regulation throughout neuronal development is crucial for determining appropriate models for TANGO testing. During neuronal development, alternative splicing of RNA transcripts is highly regulated^30,31^. In mice, switching in splicing during neuronal maturation is regulated by neuronal RNA-binding proteins including NOVA, RBFOX, MBNL, and PTBP^30^. In hiPSC-derived neurons, it was shown that PTBP1 and PTBP2 binding to *SYNGAP1* mRNA promotes A3SS inclusion and subsequent nonsense-mediated decay^14,16^. However, data from mice show that Ptbp controls a splicing regulatory program specific to early neuronal maturation by repressing expression of adult protein isoforms^32^, and Ptbp levels decrease postnatally^33^. For therapy development, more knowledge on PTBP1 and PTBP2 levels and alternative splicing in the postnatal human brain is crucial.

Previous reports did show an increase in SYNGAP1 protein levels after treatment with SSOs in neuronal cultures^16^ and cerebral organoids^14^. In these types of long-term cultures, neurons are relatively mature. However, it is important to realize that both these models are not neuronal monocultures, but also contain non-neuronal cell types like astrocytes and neuronal progenitors. Therefore, it is unclear if the increases in SYNGAP1 protein levels originate from the neurons, or the other cell types present in the culture. Because the A3SS is the predominant isoform in non-neuronal cell types, while it is low in neurons, we deem it more likely that the protein resulting from SSO-mediated isoform switching originates from non-neuronal cell types. In this case, it is unlikely that the SSOs functionally rescue the *SYNGAP1* haploinsufficiency phenotype. However, it is possible that a small increase in SYNGAP1 protein levels could already give clinical benefit. Electrophysiological studies assessing the neuronal function of SSO-treated cultures would give more clarity on the SSO’s therapeutic potential.

We showed that *SYNGAP1* haploinsufficient NGN2 neurons show earlier maturation of action potential properties compared to isogenic controls, which is in line with findings that *SYNGAP1* haploinsufficiency disrupts neoteny^34^. At week 3, action potential amplitude was higher in mutant neurons compared to isogenic controls, in line with recent findings in hiPSC-derived neurons harboring a c.435_447dup mutation^35^. Because *SYNGAP1* haploinsufficiency leads to a clear electrophysiological phenotype, this could be used as a read-out to assess the effect of potential treatments. However, caution is warranted: we only observed these differences at week 3 and not at week 5, meaning this phenotype could be related to neuronal maturity. More research into the translatability of the electrophysiological findings in hiPSC-derived models compared to the human condition is recommended.

Together, our findings highlight the importance of taking the maturity and cellular heterogeneity of hiPSC-derived models into account for testing therapeutic approaches based on splice-switching for *SYNGAP1* haploinsufficiency and other neurodevelopmental disorders.

## Materials and Methods

### Cell line generation

The parental iPSC line was derived from fibroblasts of a SYNGAP1-related disorder patient (GM27957, female, 4 years)^22^ harboring a heterozygous *de novo* loss-of-function mutation at Exon 17 (c.3718 C>T). The patient mutation in the parental iPSC line was corrected using CRISPR/Cas9 genome editing to create an isogenic control line; a clonal line that was carried through the same workflow but was not edited at the *SYNGAP1* locus and therefore retained the patient mutation was used as the *SYNGAP1* mutant line. CRISPR/Cas9 genome editing was performed by delivering a ribonucleotide protein (RNP) consisting of S.p.Cas9-RFP V3 (IDT, 10008163), and a guide RNA targeting the patient allele (GATGCAGTATCAGGCCTGAC, IDT). The RNP was delivered together with a single-stranded oligonucleotide with Alt-R modifications (GAGCGGAGGCTGCTGTCCCAGGAAGAACAAACCAGCAAAATCCTGATGCAGTATCA GGCCCGACTAGAGCAGAGTGAGAAGAGGCTAAGGCAGCAGCAGGCAGAGAAGGATTCCCAGATCAAGAGC, IDT) using the P3 Primary Cell 4D-Nucleofector X Kit (Lonza, V4XP-3024) and nucleofected using program CA-137 of the Lonza 4D-Nucleofector X Unit. Control and mutant iPSC lines were used to create Neurogenin 2 (NGN2) iPSCs for neuronal differentiation. Cells were nucleofected with S.p.Cas9-RFP V3 (IDT, 10008163), and guide RNA targeting *AAVS1* (GGGGCCACTAGGGACAGGAT, IDT) and a donor plasmid (Addgene #105840 with mCherry swapped for eGFP as a fluorescence marker) to integrate the NGN2 construct containing a puromycin-resistant gene at the *AAVS1* locus. Nucleofection was performed using the P3 Primary Cell 4D-Nucleofector X Kit (Lonza, V4XP-3024) and program CA-139 of the Lonza 4D-Nucleofector X Unit. Transfected colonies were selected by addition of puromycin (300 ng/ml, ThermoFisher Scientific, A1113803) to the iPSC medium.

### hiPSC culture

hiPSCs were cultured in 6-well plates coated with Geltrex (Life Technologies, A1413302). For passaging, cells were dissociated with Accutase (A1110501, ThermoFisher Scientific), centrifuged in Dulbecco’s phosphate buffered saline (DPBS; 14190169, Life Technologies) at 1000 x g for 3 min, and resuspended and plated in iPSC medium (Table 1). Medium was refreshed with iPSC medium without RevitaCell (ThermoFisher Scientific, 15317447) the day following thawing and passaging and every two or three days after that. Cells were cultured at 37 °C and 5% CO_2_. All hiPSC lines and their derivatives underwent regular microarray-based screening for structural genomic variation and were screened for mycoplasma every other month. Because the parental line was female, we additionally monitored X chromosome inactivation (XCI) status at early passages. The line showed evidence of partial XCI erosion at baseline. All experiments were performed at early passage numbers to limit further erosion.

**Table 1:** Cell Culture Media.

| Name | Reagents | Manufacturer, Catalogue Number |
| --- | --- | --- |
| iPSC Medium | StemFlex medium | ThermoFisher Scientific, A3349401 |
|  | 1% Penicillin-Streptomycin | ThermoFisher Scientific, 15140122 |
|  | 1% RevitaCell | ThermoFisher Scientific, 15317447 |
| Neural | Advanced DMEM/F12 | ThermoFisher Scientific, 1634010 |
| Induction | 1% Penicillin-Streptomycin | ThermoFisher Scientific, 15140122 |
| Medium | 1% N-2 Supplement | ThermoFisher Scientific, 17502048 |
|  | 2 ug/ml heparin | Merck, H3149-10KU |
|  | 20 ng/ml basic FGF | Merck, GF003AF |
| NPC Medium | Advanced DMEM/F12 | ThermoFisher Scientific, 1634010 |
|  | 1% Penicillin-Streptomycin | ThermoFisher Scientific, 15140122 |
|  | 1 ug/ml Laminin | Sigma-Aldrich, L2020 |
|  | 1% N-2 Supplement | ThermoFisher Scientific, 17502048 |
|  | 2% B-27 Minus RA Supplement | ThermoFisher Scientific, 12587010 |
|  | 20 ng/ml basic FGF (for NPCs) | Merck, GF003AF |
|  | 10 ng/ml BMP4 (for Astrocytes) | ProSpec, CYT-361 |
|  | 10 ng/ml LIF (for Astrocytes) | PeproTech®, 300-05-25UG |
| Day 1 Medium | Advanced DMEM/F12 | ThermoFisher Scientific, 1634010 |
|  | 1% Penicillin-Streptomycin | ThermoFisher Scientific, 15140122 |
|  | 1% MEM-NEAA | ThermoFisher Scientific, 11140035 |
|  | 1% N-2 Supplement | ThermoFisher Scientific, 17502048 |
|  | 4 ug/ml doxycycline | ThermoFisher Scientific, 12587010 |
|  | 10 ng/ml NT3 | PeproTech, 450-03 |
|  | 20 ng/ml BDNF | ProSpec, CYT-207 |
|  | 200 ng/ml Laminin | Sigma-Aldrich, L2020 |
| NGN2 | Neurobasal Medium | ThermoFisher Scientific, 21103049 |
| Medium | 1% Penicillin-Streptomycin | ThermoFisher Scientific, 15140122 |
|  | 1% Glutamax | ThermoFisher Scientific, 35050061 |
|  | 2% B-27 Minus RA | ThermoFisher Scientific, 12587010 |
|  | 4 ug/ml doxycycline | ThermoFisher Scientific, 12587010 |
|  | 10 ng/ml NT3 | PeproTech, 450-03 |
|  | 10 ng/ml BDNF | ProSpec, CYT-207 |
|  | 2 ng/ml T3 | Sigma Aldrich, 642511 |

### Generation of neural progenitor cells

iPSC lines were used to generate NPCs through Embroid Body (EB) differentiation in neural induction medium^23^ (**Table 1**). After 7 days in suspension, EBs were plated on laminin coated dishes (20 ug/ml; Sigma, L2020). From day 14 onwards, plated EBs were refreshed with NPC medium (**Table 1**). Cells were passaged 1:4 every week using collagenase (ThermoFisher Scientific, 17104019) At passage 3, NPCs were purified by fluorescence-activated cell sorting (FACS)^36,37^. Cells were detached using Accutase^TM^ cell dissociation reagent (ThermoFisher Scientific, A1110501) and resuspended in a single cell solution in 2% FBS (ThermoFisher Scientific, 10082147) in PBS. CD184+/CD44-/CD271-/CD24+ cells were collected using a FACSAria III (BD Bioscience).

### Generation of astrocytes

NPCs were differentiated into astrocytes on laminin coated dishes (20 ug/ml; Sigma-Aldrich, L2020) by supplementing NPC medium with BMP4 and LIF(**Table 1**) as described previously^29^. Cells were passaged by detaching the cells with Accutase^TM^ and cultured at 37 °C and 5% CO_2_.

### Neuronal differentiation

The protocol to differentiate iPSCs into NGN2 neurons was adapted from Frega *et al.* (2017)^38^. Briefly, iPSCs were plated on coverslips coated with Geltrex in 12-well plates in iPSC medium containing 4 µg/ml doxycycline (Table 1). The following day, medium was refreshed with day 1 medium (Table 1). The next day, astrocytes were added to NGN2 cultures at a ratio of 1:1 to support neuronal maturation^29^. Astrocytes were generated from NPCs from a WTC iPSC line (WTC-11; #GM25256, RRID:CVCL_Y803) according to the protocol previously described by Lendemeijer *et al.,* 2024^29^. No astrocytes were added to NGN2 cultures harvested for Western blot experiments. All medium was replaced by NGN2 medium the following day. From DIV5 onwards, half the medium was refreshed every other day with NGN2 medium (**Table 1**). Cytarabine (2 µM) was added on day 3 to eliminate replicating cells. Doxycycline was no longer added to the medium after DIV14.

### Immunocytochemistry and Confocal Microscopy

Cell cultures on coverslips were fixed in 4% formaldehyde in PBS for 15 minutes. Primary antibodies (**Table 2**) were incubated overnight at 4°C in a labelling buffer containing 0.05 M Tris, 0.9% NaCl, 0.25% gelatin, and 0.5% Triton X-100 (pH 7.4). Secondary antibodies (Alexa-488, Alexa-647, and Cy3 (Jackson ImmunoResearch) were incubated for 1 hour at room temperature in the labelling buffer. Coverslips were mounted in Mowiol 4-88 (Fluka) on microscope slides. Slides were imaged with a Zeiss LSM 800 confocal microscope using ZEN software (Zeiss).

**Table 2:**
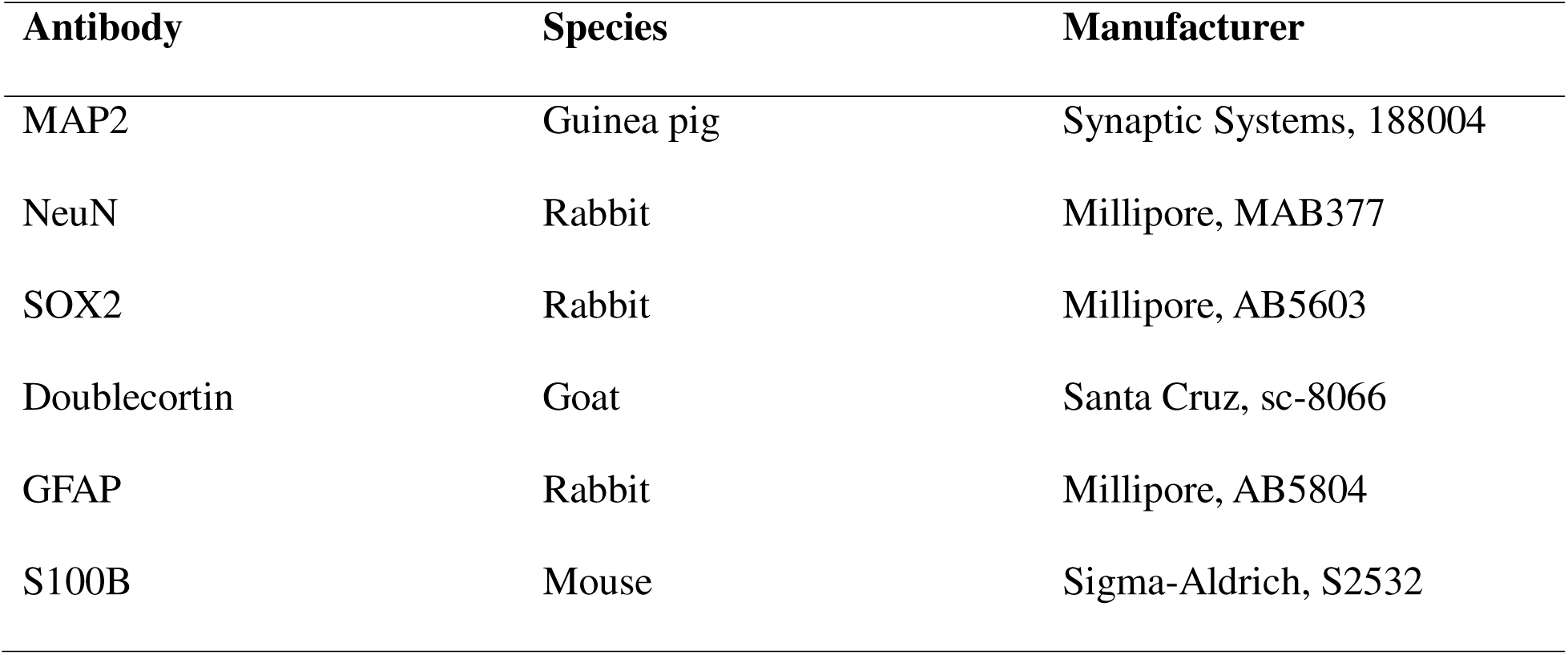
Primary antibodies.

### Whole-cell patch clamp electrophysiology

Whole cell patch clamp recordings of two batches of patient-derived and isogenic control NGN2 neurons were performed at DIV 21-25 and DIV 37-39. The recording chamber was kept at 95% O□ and 5% CO□ and perfused with 1-2 ml/min artificial cerebrospinal fluid composed of 140 mM NaCl, 2.4 mM KCl, 2 mM CaCl□, 2 mM MgCl□, 10 mM glucose, and 10 mM HEPES (pH 7.2–7.3). Temperature was regulated using a heat block set at 36-38 °C to approximate physiological conditions. Neurons were visualized using either a Nikon microscope with infrared illumination and differential interference contrast, or with a CCD camera (coolSNAP EZ, Photometrics) using Prairie View imaging software (Bruker). Patch electrodes (3–6 MΩ) were filled with intracellular solution consisting of 125 mM K-gluconate, 15 mM NaCl, 10 mM HEPES, 0.2 mM EGTA, 2 mM MgATP, 0.3 mM NaGTP, and 10 mM K phosphocreatine (pH 7.2–7.4), supplemented with 0.5% biocytin for morphological analyses. Whole-cell recordings were acquired with multiclamp 700B amplifiers (Axon Instruments), using a low-pass filter of 4 Hz and digitized at 20 kHz using Digidata 1440A (Molecular Devices) acquisition interfaces for batch 1. For batch 2, recording were acquired with HEKA EPC10 quattro amplifiers, using a low-pass Bessel filter at 2.9 kHz and digitized at 10 or 20 kHz using Patchmaster Software.

In current-clamping experiments, membrane potential was kept at -60 mV, while 500 ms currents with a frequency of 0.2 Hz were applied in 5 pA increments from the holding current. Series resistance and pipette capacitance were continuously monitored and adjusted using bridge and capacitance compensation. The action potentials were analyzed using AxoGraph with + 10 mV amplitude threshold. For voltage-clamp set-ups, pipette capacitance was compensated and recordings with a series resistance lower than 20 MΩ were included in analyses. Spontaneous excitatory postsynaptic currents (sEPSCs) were recorded for 3-5 min using a gap-free acquisition mode while cells were clamped at -60 mV and detected with Mini analysis software (Synaptosoft Inc) and Igor Pro using a Neuromatics plugin. Membrane capacitance was measured at the beginning of each recording by applying a 5 mV voltage step from a holding potential of -60 mV using the membrane test function in Clampex or Patchmaster.

### Western blot

Cultured NPCs, astrocytes, and NGN2 neurons were harvested in 1% protease inhibitor cocktail (Sigma-Aldrich, P8340) *in triplo*. Samples and reused mouse brain lysates as positive controls were homogenized in Laemmli lysis buffer (1M Tris-HCl at pH 7.5, 4% SDS). Protein concentrations were measured photometrically using BCA protein assay kit (Pierce^TM^, Thermo-Fisher Scientific). Samples were diluted in 25% XT Sample Buffer (BioRad) containing 2.5% β-mercaptoethanol and supplemented with water before gel electrophoresis. Samples were heat-shocked at 70-95°C for 5-10 minutes and electrophoresed through 4-12% Criterion^TM^ XT Bis-Tris gradient gels (BioRad) in XT-MOPS Buffer (BioRad). Proteins were transferred to 0.45 um pore nitrocellulose membranes (BioRad) in a buffer consisting of 10% Tris-Glycine (BioRad) and 20% anhydrous methanol in distilled water. Membranes were blocked in TBS-T buffer containing 0.1% Tween 20 (Sigma Aldrich, P1379) and 5% Blotting Grade Blocker (Bio-Rad, 1706404XTU) at room temperature for 2 hours. Primary antibodies against SYNGAP1 (Invitrogen, PA1 046) and glyceraldehyde 3-phosphate dehydrogenase (GAPDH; Cell Signaling Technology, 14C10) or Actin (Millipore, MAB1501R) were incubated overnight at 4 °C in TBS-T with 2% Blotting Grade Blocker. After 3x5 min washes in TBS-T, membranes were incubated with IRDye secondary antibodies (LICORbio) in TBS-T for 45 min at room temperature and subsequently washed 3x5 min with TBS-T and 1x5 min with TBS. Membranes were imaged on an Odyssey M system (LICORbio). Gel band intensities were quantified using ImageJ.

### RNA sequencing for alternative splicing analysis

To assess *SYNGAP1* splicing across different hiPSC-derived brain cell types, we interrogated a previously generated NMD-sensitive alternative splicing (NMD-AS) database containing long-read and short-read RNA sequencing data from hiPSCs, *ngn2*-induced neurons (DIV49), iPSC-derived astrocytes (DIV42), and iPSC-derived microglia (DIV23)^22^. Because of insufficient long-read sequencing coverage, *SYNGAP1* alternative splicing events were excluded from the filtered long-read dataset, and therefore the unfiltered long-read dataset, in which these events were retained, was used to identify *SYNGAP1* splicing events. This revealed the previously described A3SS event (event ID: A3:6:33438919-33440553:33438919-33440729:+). Percent spliced-in (PSI) values for this event were calculated from the short-read RNA sequencing data using the three biological replicates available for each cell type and treatment.

### Reverse Transcription (RT)-PCR

Cultured NPCs, astrocytes, and NGN2 neurons were harvested *in triplo* for Reverse Transcriptase PCR (RT-PCR) after being treated with cycloheximide (200 ug/ml; Sigma-Aldrich, CAS66-81-9) for 5 hours to inhibit NMD. NGN2 neurons that were co-cultured with control astrocytes were purified by GFP using FACSAria III (BD Bioscience) before RNA extraction. RNA extraction was performed with RNeasy Mini Kit (Qiagen, 74104). cDNA was synthesized according to instructions of the iScript cDNA Synthesis Kit (Bio-Rad, 1708891). DNA around the A3SS inclusion was amplified with the primer set used by Yang et al. (2023; forward: 5’-CAAGGATGCCATTGGAGAAT-3’; reverse 5’-GCAAACACCTCCTTCAGCTC-3’). The PCR program consisted of an initial denaturation step of 94 °C for 3 min, followed by 45 cycles of denaturation at 94 °C for 45 seconds, annealing at 52 °C for 1 min and extension at 72 °C for 2 min, and a final extension step at 72 °C of 10 min. DNA was visualized with SYBR Green PCR Master Mix (Thermo Fisher Scientific, 4309155) and loaded on a 1.5% agarose gel, together with a FastRuler Low Range ladder (ThermoFisher Scientific, SM1103). Gel band intensities were quantified using ImageJ.

### Statistics

Before statistical analyses, data distributions were checked with Q-Q plots, histograms and Shapiro-Wilk tests for normality. Electrophysiological measurements were statistically compared with linear mixed models including genotype and timepoint as fixed and batch as random effect. Log-transformed values were fitted for properties that were not normally distributed. Post-hoc comparisons were conducted using estimated marginal means and p-values were adjusted with a False Discovery Rate correction. Corrected p-values lower than 0.05 were considered significant. For Western blot experiments in NGN2 neurons, SYNGAP1/GAPDH ratios were statistically compared using a two-way ANOVA model and post hoc Tukey test. For RT-PCR experiments in NGN2 neurons, mean PSI scores using a two-way ANOVA model and p-values were adjusted with an FDR correction.

## Acknowledgements

This work was supported by the ZonMw PSIDER program TAILORED (10250022110002) and by European Union’s Horizon Europe Research and Innovation Programme ‘EURAS’ (101080580).

## Author contributions

J.A.K., S.A.K. and F.M.S.d.V. developed the concept for the paper. J.A.K and D.G. generated the CRISPR/Cas9-edited hiPSC lines. M.A.D.J and N.M. performed the patch clamping and analyzed the raw data. C.G. generated the NGN2 plasmid and performed XCI checks. K.N.W. performed the RNA sequencing and analyzed the data. J.A.K., D.G., and M.R. performed the remaining experiments and analyzed the data. J.A.K. and F.M.S.d.V. wrote the manuscript with input from all co-authors. J.A.K., S.A.K, Y.E., L.E.L.M.V., N.N.K and F.M.S.d.V. supervised the work.

