## Supplementary material for "Maturation-dependent splicing alterations constrain *SYNGAP1* splice-switching therapy": Suplemental table 1

**Supplemental table S1 Descriptives of electrophysiological properties of control and *SYNGAP1* mutant NGN2 neurons at weeks 3 and 5.** Values are represented as mean±standard deviation or median (interquartile range).

| **Property** | **Control W3** | **Mutant W3** | **Control W5** | **Mutant W5** |
| --- | --- | --- | --- | --- |
|  | *n = 26* | *n = 30* | *n = 26* | *n = 26* |
| Action potential half-width (ms) | 1.82 (0.57) | 1.32 (0.35) | 1.24 (0.28) | 1.27 (0.46) |
|  | *n = 16* | *n = 21* | *n = 26* | *n = 26* |
| Action potential rise time (ms) | 0.56 (0.22) | 0.38 (0.11) | 0.36 (0.08) | 0.39 (0.14) |
| Action potential decay time (ms) | 0.98 (0.33) | 0.77 (0.19) | 0.70 (0.16) | 0.69 (0.28) |
|  | *n = 30* | *n = 32* | *n = 26* | *n = 26* |
| Action potential peak voltage (mV) | 62.39±17.90 | 80.14±17.81 | 81.98±8.65 | 77.70±12.99 |
| Action potential threshold (mV) | -19.83±12.79 | -21.43±8.11 | -21.34±7.02 | -21.32±6.72 |
|  | *n = 34* | *n = 36* | *n = 26* | *n = 31* |
| Membrane capacitance (pF) | 27.06±13.00 | 30.93±14.53 | 62.81±22.05 | 62.39±15.17 |
| Resting membrane potential (mV) | -37.03±10.35 | -40.46±7.40 | -45.12±5.07 | -43.87±8.94 |
| Maximum firing rate (Hz) | 7.84±5.45 | 10.19±4.19 | 13.46±2.58 | 12.69±3.72 |
| Membrane resistance (MΩ) | 495 (330) | 685 (665) | 420 (302.5) | 500 (455) |
| Input resistance (MΩ) | 769.64 (629.08) | 1020.16 (625.53) | 415.48 (364.10) | 536.54 (376.85) |
|  | *n=18* | *n=22* | *n=26* | *n=31* |
| sEPSCs frequency | 0.005 (0.019) | 0.072 (0.130) | 0.967 (1.127) | 0.777 (0.543) |
