## Supplemental figure 1 for "Maturation-dependent splicing alterations constrain *SYNGAP1* splice-switching therapy"

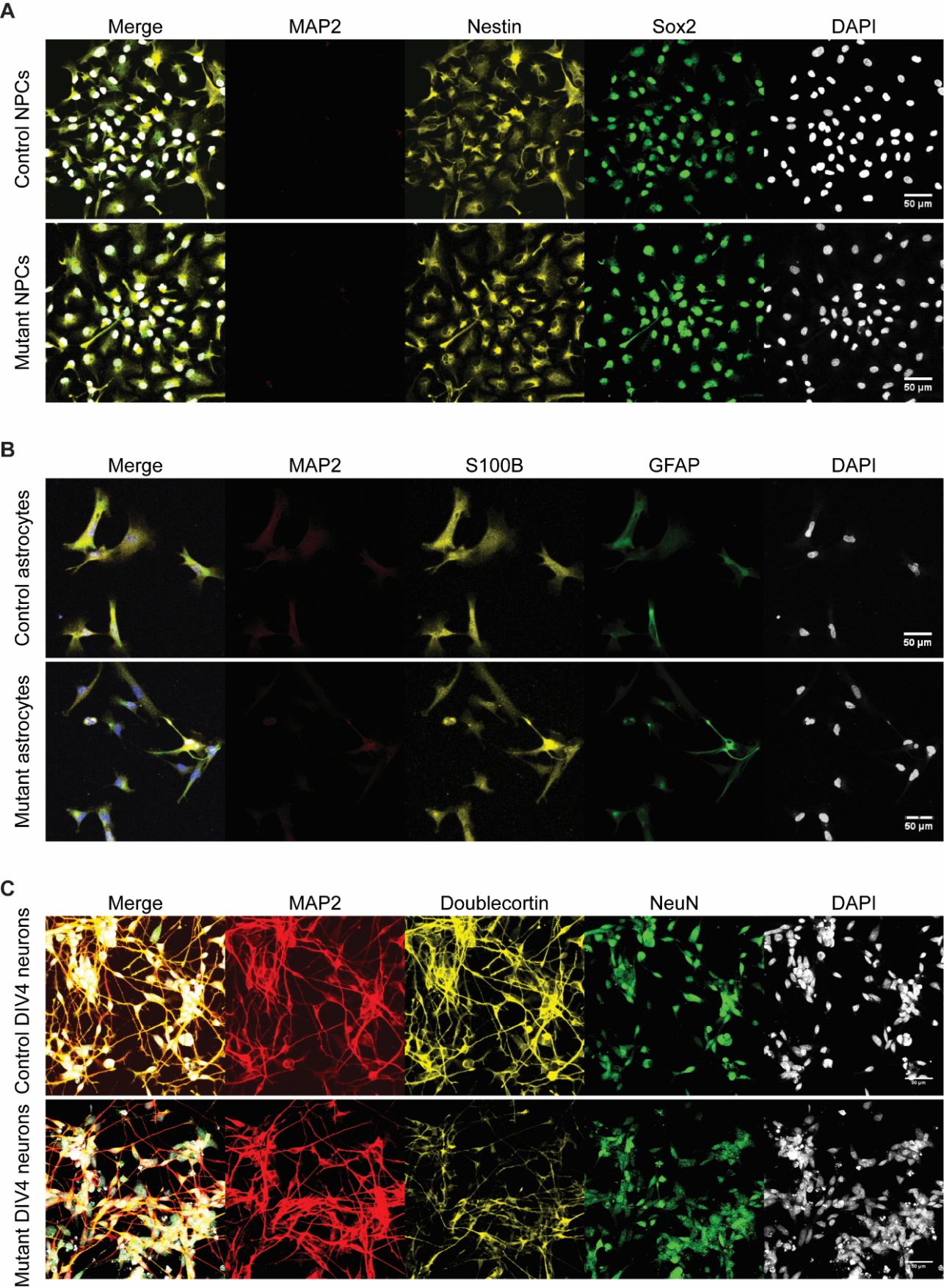

**Figure S1 SYNGAP1 mutant and isogenic control neural progenitors, astrocytes and DIV4 neurons used in this study express the expected markers** Representative pictures of (A) neural progenitors stained for MAP2, Nestin, Sox2 and DAPI (B) astrocytes stained for MAP2, S100B, GFAP and DAPI and (C) DIV4 neurons stained for MAP2, Doublecortin, NeuN and DAPI. Scale bars depict 50 μm.
